# Engineered AAV Capsids with Enhanced Extracellular Vesicle Loading via Rational Design and Directed Evolution

**DOI:** 10.64898/2025.12.12.693981

**Authors:** Ioulia Vorobieva, Marina Luchner, Hans Friedrichsen, Ruhani Makkar, Simon Chester, Emily Haughton, Mariana Conceiao, Harrison Steel, Stephan J. Sanders, Matthew PA Wood, Dhanu Gupta

**Author notes:** These authors contributed equally to the manuscript (Ioulia Vorobieva, Marina Luchner).

## Abstract

Overcoming immune responses and ensuring efficient delivery remain key obstacles to advancing adeno-associated virus (AAV)-based genetic medicines into the clinic. Encapsulating AAVs within naturally secreted extracellular vesicles (EV-AAV) can shield them from neutralising antibodies and increase transduction efficiency; however, wild-type AAV capsids are incorporated into EVs inefficiently, with the percentage of total AAV genomes in the EV-AAV fraction ranging from 0.5% to 12%, depending on AAV serotype. Here, we establish a high-throughput screening platform that links AAV capsid genotype to EV encapsulation phenotype to improve encapsulation efficiency. We combined rational design with directed evolution to engineer AAV9 variants that preferentially sort into EVs. A library of 392,550 AAV9 capsid mutants was generated by inserting Late (L) domains, established EV-sorting motifs, and fully random amino acid inserts in a surface-exposed region of the AAV9 capsid protein. Screening data were analysed using a statistical model-assisted protein selection (SMAPS) workflow – developed as a novel quantitative modelling and selection strategy for protein engineering challenges of this kind. SMAPS functions as a digital twin of the protein variants in our high-throughput screen, allowing us to monitor uncertainty in copy numbers over multiple steps in the experimental workflow. Using SMAPS, we identified five lead hits, which showed up to threefold increased EV-AAV yield compared to wild-type, without altering particle size. Moreover, *in vitro* transduction experiments demonstrated that the lead AAV9 capsid exhibits substantially higher transduction efficiency compared to wild-type AAV9 and retained functional activity in the presence of neutralising anti-AAV9 antibodies. Collectively, our study: (i) delivers a scalable high-throughput screen for EV-AAV engineering, (ii) introduces the field’s first statistical model-guided quality-control framework for directed evolution, and (iii) provides best-in-class capsids for EV encapsulation.

## Introduction

Adeno-associated virus (AAV) vectors are widely used in genetic medicines due to their relatively favourable safety profile, sustained transgene expression, broad tissue tropism and limited integration of DNA into the genome, with seven AAV-derived therapeutics having received regulatory approval so far^1–3^. However, the clinical efficacy of AAV-based therapies remains limited by several key challenges, including pre-existing immunity, inefficient delivery to target tissues, manufacturing bottlenecks, and liver toxicity^4–6^. One emerging strategy to overcome some of these hurdles involves the use of extracellular vesicle-associated AAVs (EV-AAVs). EVs are lipid-bound nanoparticles naturally released by cells that are capable of mediating intercellular communication by transferring cargo such as proteins and nucleic acids^7^. Several non-enveloped viruses—including hepatitis A virus and poliovirus, as well as rotavirus and norovirus—are released inside extracellular vesicles that cloak virions, enhance infectivity, and confer partial resistance to neutralization. Similarly, AAVs within or associated with EVs have been shown to enhance cellular uptake and evade host immune detection ^9–12^.

Despite these advantages, wild-type AAVs are inefficiently loaded into EVs during production, with serotypes AAV2, AAV8, and AAV9 showing that only 2–4% of secreted viral genomes associate with EVs, and only 1.2–1.6% of total EVs secreted during production incorporating AAVs (SupFig1). Further, the percentage of total AAV genomes in the EV-AAV fraction ranges from 0.5% to 12%, depending on serotype^13^. To address this, rational engineering of the AAV capsid to incorporate cellular trafficking motifs offers a promising strategy to improve EV association. In particular, viral late assembly domains (L domains) and EV-sorting signals such as KFERQ have been identified in other viruses as mediators of interactions with the host cell machinery involved in vesicle formation, such as the ESCRT (endosomal sorting complexes required for transport) pathway^14–16^. Incorporating such motifs into the AAV capsid may promote packaging into EVs by hijacking these endogenous vesicle biogenesis pathways. However, the incorporation of such motifs is limited by the structural constraints of the capsid, as larger insertions can disrupt capsid assembly, stability and folding, leading to reduced AAV yield or functionality^17^.

Another approach to enhance the EV encapsulation efficiency of AAVs is directed evolution, which has been successfully adopted to engineer AAV variants with enhanced tissue tropism^18^ and immune evasion properties^19^. Directed evolution experiments sample the protein fitness landscape via trial-and-error by mutating the gene encoding the protein of interest and selecting for high-performing variants, leading to the accumulation of variants with enhanced properties^20^. As opposed to rational design, detailed understanding of the underlying biophysical mechanisms is not required for directed evolution. This is particularly interesting for our engineering problem since the EV encapsulation mechanisms of AAVs are (so far) not well understood.

In this study, we employed a combination of rational design and directed evolution to engineer AAV9 capsids with enhanced EV incorporation. We generated a library of 392,550 AAV9 capsid variants by inserting L domains, established EV-sorting motifs or randomised sequences, into a surface-exposed loop of the AAV9 capsid. The library was screened across multiple rounds of EV enrichment and high-throughput sequencing to identify capsid variants that exhibited superior association with EVs. Alongside experimental screening, we developed a statistical model-assisted protein variant selection strategy (SMAPS) to support high-throughput analysis and enrichment-based variant selection. The resulting platform offers both a set of capsid variants with 3-fold enhanced EV incorporation, and a generalizable analytical strategy that can be adapted by others to optimise library screening workflows. Selected variants from this platform were further evaluated in both *in vitro* and *in vivo*, validating the screen’s findings by confirming enhanced functional performance. These findings underscore the potential of capsid engineering to improve the therapeutic efficacy and manufacturability of EV-AAVs.

## Results

### A rationally designed AAV capsid library enables high-throughput screening of variants with enhanced EV loading properties

To enhance AAV capsid incorporation into EVs, we focused on engineering the surface-exposed variable region IV (VR-IV) loop of the AAV9 capsid protein. We hypothesized that the introduction of short peptide motifs known to interact with the ESCRT machinery could promote EV association. Furthermore, we reasoned that the length and flexibility of linkers flanking these motifs might modulate their accessibility and function, particularly given that this capsid insertion site has previously accommodated other peptides and is positioned to enable intracellular interactions with EV-sorting proteins. A diverse capsid library was thus designed to encode various combinations of rationally selected late assembly domains (L domains) and EV loading motifs, as well as randomised sequences of equal length (Figure 1, Table S1). Each central motif was flanked by linkers composed of either two, four, or eight random amino acids to screen for the ideal spacer length and sequence, and the constructs were inserted at residue 453 (VP1 numbering) of the AAV9 capsid.

**Figure 1.**
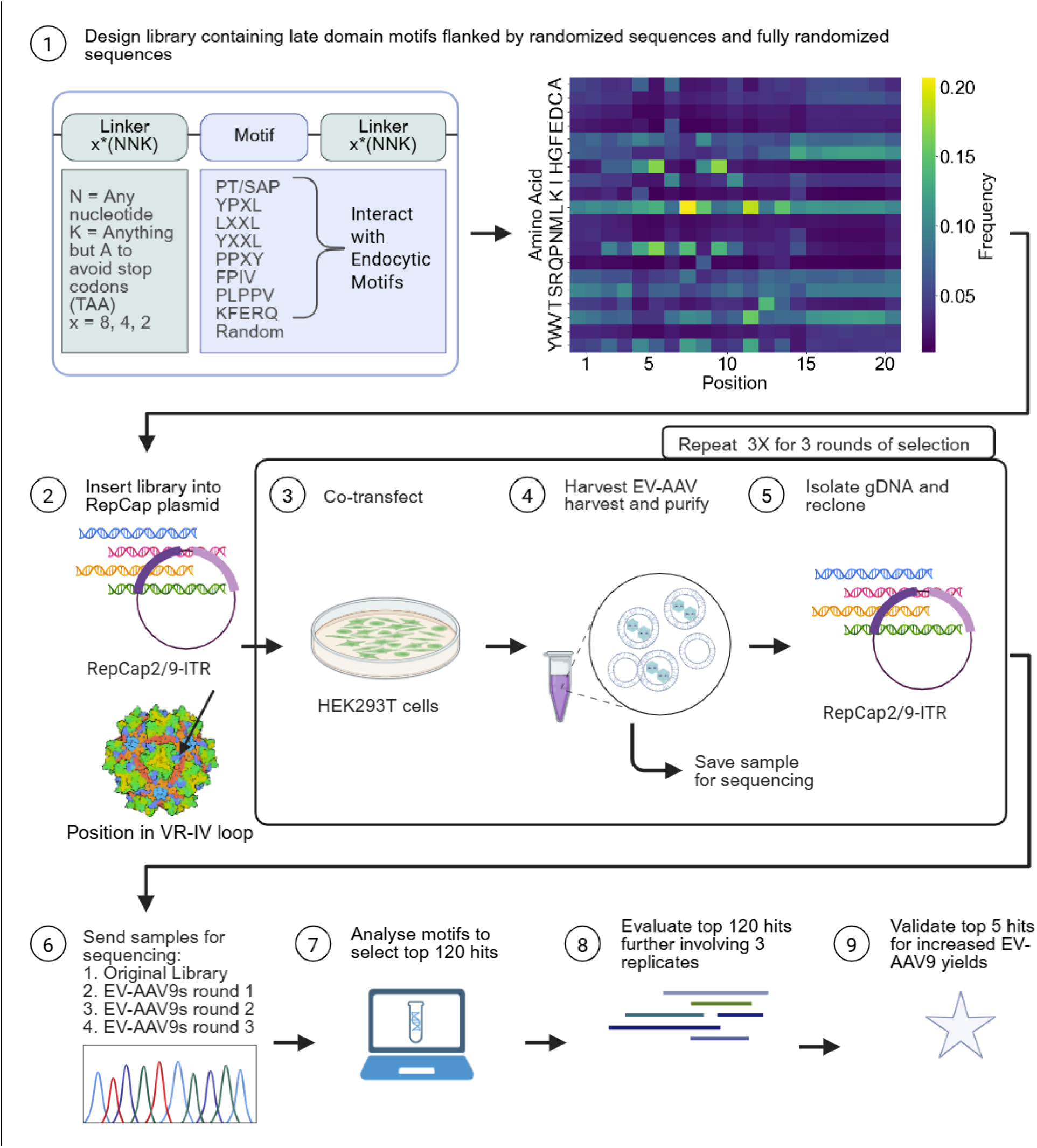
Generation of an AAV9 library to identify variants with improved extracellular vesicle incorporation efficiency. This figure illustrates the experimental workflow for creating a library of AAV9 variants and selecting for those with enhanced EV incorporation (A). First, a library was designed with randomised linkers of 2, 4, or 8 codons with either non-random amino acids encoding L domain motifs or random amino acids in the centre. The amino acid frequency at each position in the insert is displayed as a heatmap (B). These sequences were cloned into the VR-IV loop of the AAV9 capsid protein. The subsequent steps involved co-transfection of HEK293T cells with the AAV RepCap2/9-ITR plasmid, followed by EV-AAV harvest and purification. DNA was extracted from EV-AAVs and cloned back into the RepCap2/9-ITR plasmid for repeated rounds of selection. Samples from each round were then prepared for sequencing by PCR amplification of the insert region. Resulting data was used to perform motif analysis to identify specific amino acid sequences or motifs that correlated with improved EV incorporation. The top 120 hits were selected and ordered as an oligo pool for further evaluation. Three biological replicates of EV-AAV harvest were performed and sent for Amplicon EZ Sequencing. These results were used to select top 5 candidates to validate for improved incorporation into EVs.

To minimise the risk of introducing stop codons, the library was synthesised using NNK codons for the variable regions. The resulting oligonucleotides were cloned into the AAV9 rep-cap expression cassette, placed between inverted terminal repeats (ITRs) to allow the capsid variant genome to be packaged. HEK293T cells were transfected with the library, and EV-AAVs were isolated from conditioned media using iodixanol gradient ultracentrifugation. Viral DNA was extracted from the EV-enriched fractions and re-cloned into the parental vector for the next round of selection. This iterative process was performed for three rounds to enrich for capsids with enhanced EV association. Next-generation sequencing (NGS) was used to measure the number of capsid variants in the original library and the enriched library. The EV encapsulation efficiency of a variant was approximated by calculating the enrichment of the variant in the EV fraction.

### SMAPS allows for the identification of high-performing variants with an enrichment score above simulated noise levels

To identify AAV capsid variants with enhanced EV loading properties, we quantified the EV encapsulation efficiency by comparing the frequency of the AAV capsid protein variants between the EV fraction of the supernatant and the original plasmid library, resulting in an enrichment score. However, this enrichment score is associated with an error term, which is accumulated over multiple experimental processing steps (e.g., random enrichment during growth, sequencing biases). Therefore, we simulated the baseline expected range of enrichments based on random chance and identified variants enriched beyond that range (Figure 2A). We show that our model approximates the copy number distribution of the protein variants screened and that an enrichment score cutoff of 10 discards 99.99% of randomly enriched variants (Figure 2B). This result emphasizes the need for our statistical model as it demonstrates that the “noise range” is non-negligible, with 99.31% of protein variants falling below the estimated noise threshold. This modelling-based approach provides a broadly applicable analytical framework that allows separation of signal from noise for others performing high-throughput enrichment-based protein engineering screens (see Code Availability).

**Figure 2.**
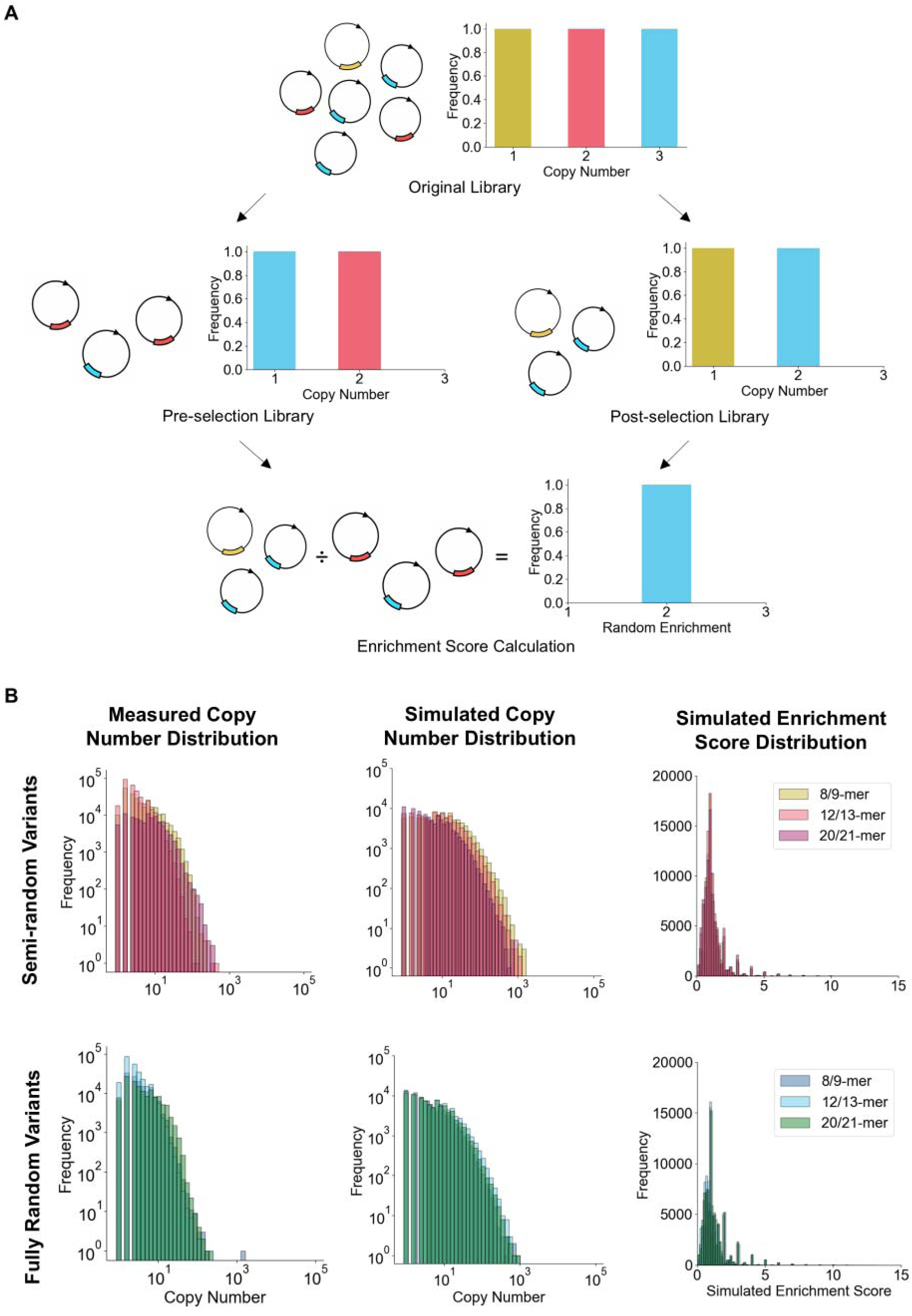
Disentangling selection through evolutionarily advantageous traits from random enrichment. We developed SMAPS, a bespoke statistical model to estimate protein variant enrichment based on random chance (A). The model describes the numerical transformations that the population of protein variants undergoes during experimental processing. As the model accounts for stochasticity, it allows for disentangling random enrichment of protein variants from enrichment through beneficial properties. We modelled experimental processing steps that subsample the library and simulated enrichment (and depletion) in the original library compared to the evolved library (B) Here, we compare between variant distributions for experimentally measured and simulated workflows. Comparing left and centre columns shows that our simulations remarkably succeed to approximate the measured copy number distribution, demonstrating the fidelity of our modelling approach. Meanwhile, simulated enrichment scores show that variants can be randomly enriched up to 10 times compared to averagely performing variants with an enrichment score of 1. This insight is crucial, as it empowers us to define a meaningful enrichment score cutoff, disentangling random enrichment from active selection.

To mitigate the risk of selecting AAV capsid variants that are enriched due to cross-packaging, we applied motif analysis, depicting amino acids as enlarged in proportion to their frequency per position. This allows for visually identifying motifs enriched across multiple variants (Figure 3A). For both the semi-random and fully-random insert designs, we picked top motifs for each insert length.

**Figure 3.**
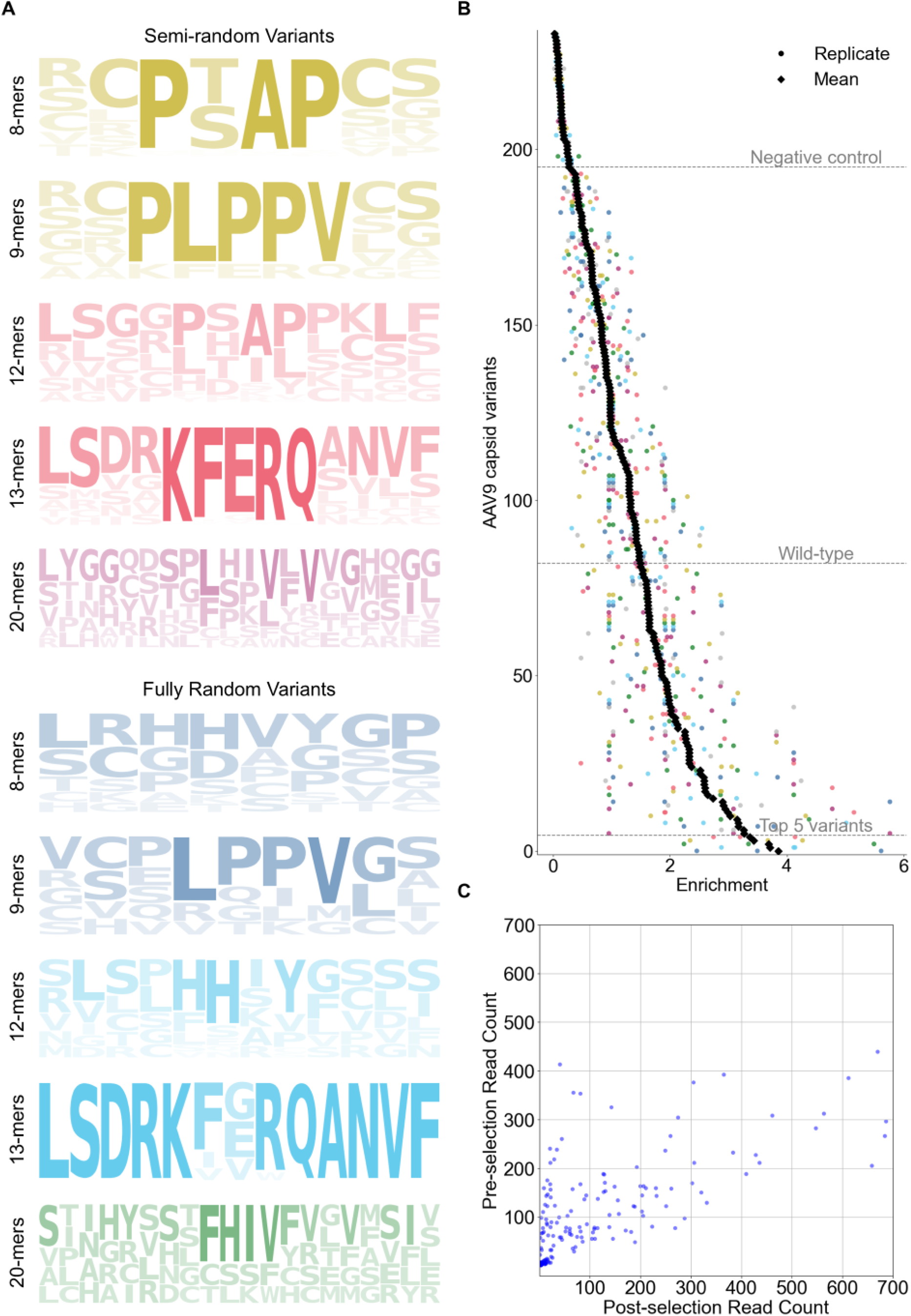
Capsid variant screening and hit selection. After completing three rounds of evolution, samples were sent for NGS. Motif analysis was applied to identify motifs that are enriched across variants (A). Top variants were shortlisted from the evolution capsid library screen and evaluated in a refined screen, which included negative controls with stop codons, while wildtype served as positive control. Enrichment scores were calculated by dividing the number of sequences post-selection by the number of sequences pre-selection and plotted for each evaluated variant (B). Notably, enrichment depends on the performance of the other variants present in the screen. Hence, enrichment scores cannot be compared directly between experiments, unless the experimental set-up is equivalent, but can be compared once normalised by wild-type performance. Read count post-selection (isolation of EV fraction) was plotted against pre-selection, displaying the range of read counts in both libraries (C). Enriched variants – winners – are found in the bottom right corner.

A total of 120 inserts containing top motifs and associated with enrichment scores above the cutoff were selected. Bioinformatically selected inserts were synthesised commercially as an oligo pool, re-cloned into the VR-IV capsid insertion site and produced in HEK293T cells across three biological replicates. EV-AAVs produced from these variants were again subjected to iodixanol purification and sequencing. We show that ∼ 30% of the shortlisted variants perform better than wild-type and are up to four times enriched compared to average AAV variants (Figure 3B-C). Amplicon sequencing of iodixanol-purified EV-AAV9 fractions identified five consistently highly enriched capsid variants: SVPPHHAYVPTG, LSGGLDILPKLC, LRGDVGGP, GRPLPPVTG, and LSDRKIERQGNVF (Table S2). These sequences contain bioinformatically selected motifs, including HH_Y, LSG, GGP, LSD and NVF, bolstering the validity of our hit selection strategy. Additionally, the presence of known or putative EV-sorting motifs, such as RGD, L L, PLPPV, and KFERQ-like elements, support a mechanistic link to EV biogenesis pathways^15,21^. These motifs have been implicated in facilitating the detachment of particles from the donor cell’s plasma membrane, endosomal sorting, and adhesion and internalisation by the target cell.

### Directed evolution and rational design produce AAV9 capsid variants with markedly enhanced extracellular vesicle loading properties

To further validate the performance of our top five EV-AAV9 variants (AAV9-V1 to AAV9-V5, corresponding to the L-domain and EV-sorting motifs identified in Figure 4A), we produced them in HEK293T cells and isolated them by iodixanol gradient (Figure 4A). Up to 3-fold higher percentage of the total AAV9 genomes were found in the media of the five engineered variants in comparison to the wildtype control (Figure 4B). Nanoflow cytometry of the media revealed no significant differences in EV particle size or concentration across variants, suggesting that capsid modifications did not alter EV biogenesis or release (Figure 4C-D). qPCR titration of iodixanol-isolated EV-AAV9s showed a 2.37-fold to 3.19-fold increase in viral genome copies with the LSDRKIEROGNVF capsid variant having the highest increase (Figure 4E). To assess capsid encapsulation, we ran an a-AAV9 ELISA, with and without Triton X-100. Without Triton X-100 only capsids which are on the EV exterior are accessible for binding, whereas detergent disrupts the vesicle membrane and releases capsids from the lumen; the increase in signal after detergent therefore reports the protected, EV-encapsulated pool. The variant SVPPHHAYVPTG demonstrated a marked increase in protected capsids: approximately 50% of particles were only detectable after Triton-X-100 permeabilization, compared to ∼10% in the wildtype control. These results suggest that engineered variants are more effectively encapsulated within vesicles rather than merely tethered to the EV surface in comparison to wild type capsids (Figure 4F).

**Figure 4.**
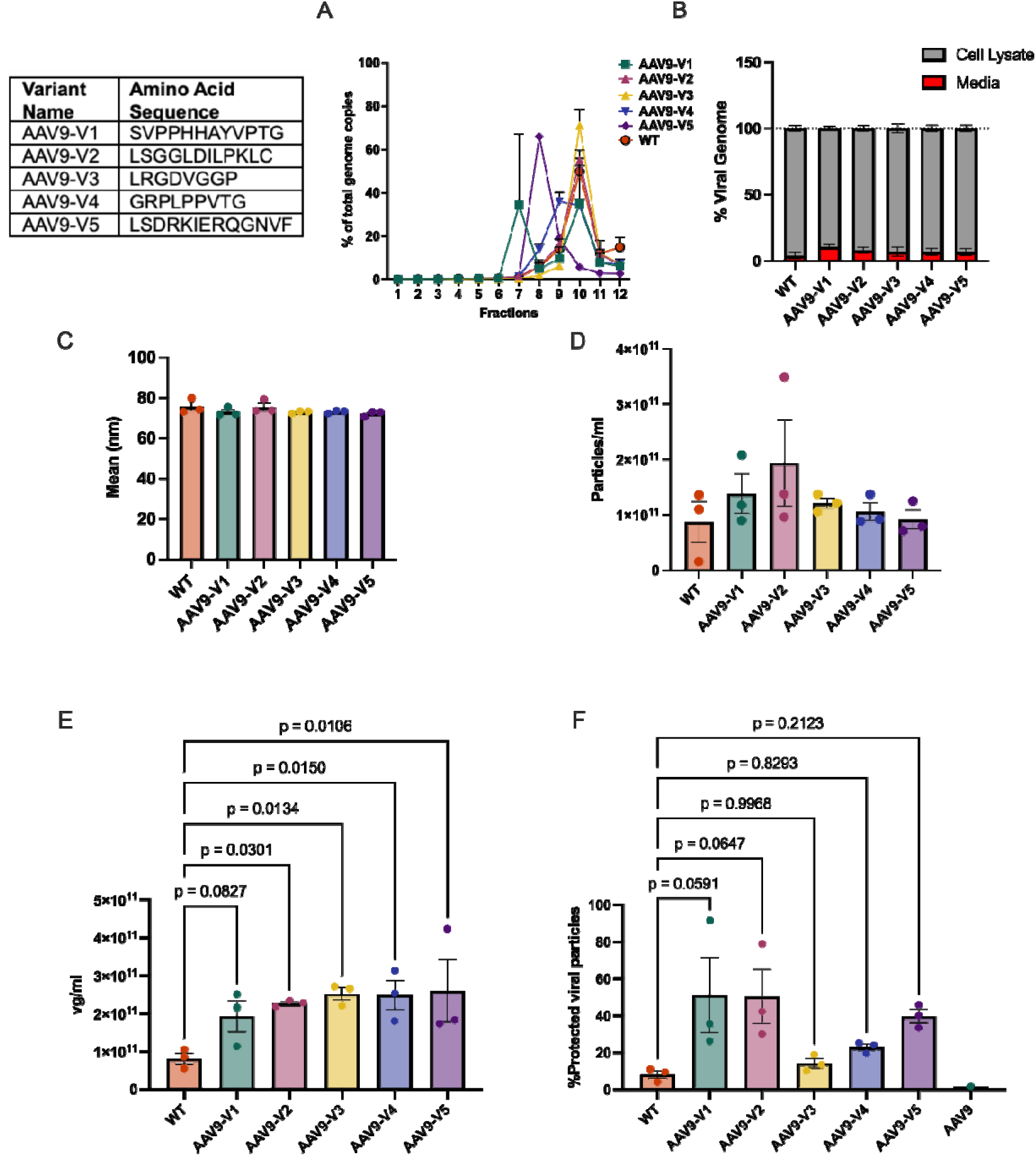
Directed evolution capsid library screen identifies capsids with increased EV-AAV9 yield. The 5 variants with the highest enrichment scores from the variant screen were selected and produced in HEK293T cells. Titration by qPCR of viral genomes in iodixanol fractions of concentrated media (A) and media and cell lysate (B). NanoFCM analysis was used to assess mean EV particle size (C) and concentration (D). qPCR titration of iodixanol-isolated EV-AAV9s samples (E). Analysis of virus particle concentration before and after treatment with Triton-X-100 (F). All values are presented as mean+SEM, N=3. Statistical analysis wa performed using GraphPad Prism, with multiple one-way ANOVA comparisons of engineered capsids to WT-EV-AAV9 control. P = Proline, T = Threonine, S = Serine, A = Alanine, Y = Tyrosine, X = Any amino acid, L = Leucine, V = Valine, F = Phenylalanine, K = Lysine, E = Glutamate, R = Arginine, Q = Glutamine.

Altogether, these findings demonstrate that our directed evolution and rational capsid design method identified AAV9 variants not only with enhanced EV association but also improved encapsulation, and transduction. This is remarkable as our screen was mainly designed to support association of AAVs with EVs and demonstrates the advantage of high throughput evolution-based screening to allow for multi-trait engineering. The enhanced properties presented here establish a strong foundation for evaluating functional performance of our lead variants in the presence of immune challenges and therapeutic delivery scenarios.

### AAV9 capsid variant 3, identified by our screening strategy has improved transduction compared to wild-type

A common challenge in directed evolution is that improvements in a selected trait may compromise other properties required for effective in vivo performance. Because our screen focused on enhancing EV loading, we first evaluated whether the engineered AAV9 variants retained intrinsic transduction capacity in vitro. All capsids carried a CMV-driven NanoLuc– eGFP fusion reporter, enabling complementary quantitative and single-cell readouts from the same vector. HeLa cells were transduced across a range of MOIs, and transduction efficiency was measured by bulk NanoLuc luminescence (Fig. 5A) and the percentage of eGFP-positive cells (Fig. 5B–C). Across all conditions, AAV9-V3 exhibited markedly enhanced transduction compared with wild-type AAV9. At the highest MOI, V3 produced nearly an order-of-magnitude increase in reporter signal, and this advantage persisted as the MOI decreased, demonstrating superior infectivity even under low-dose conditions. Flow cytometry corroborated these findings, with V3 yielding the highest proportion of eGFP-positive cells at every MOI. By contrast, AAV9-V4 and AAV9-V5 showed reduced performance—particularly at lower MOIs— indicating functional trade-offs introduced by their respective mutations that were not observed in V3.

**Figure 5.**
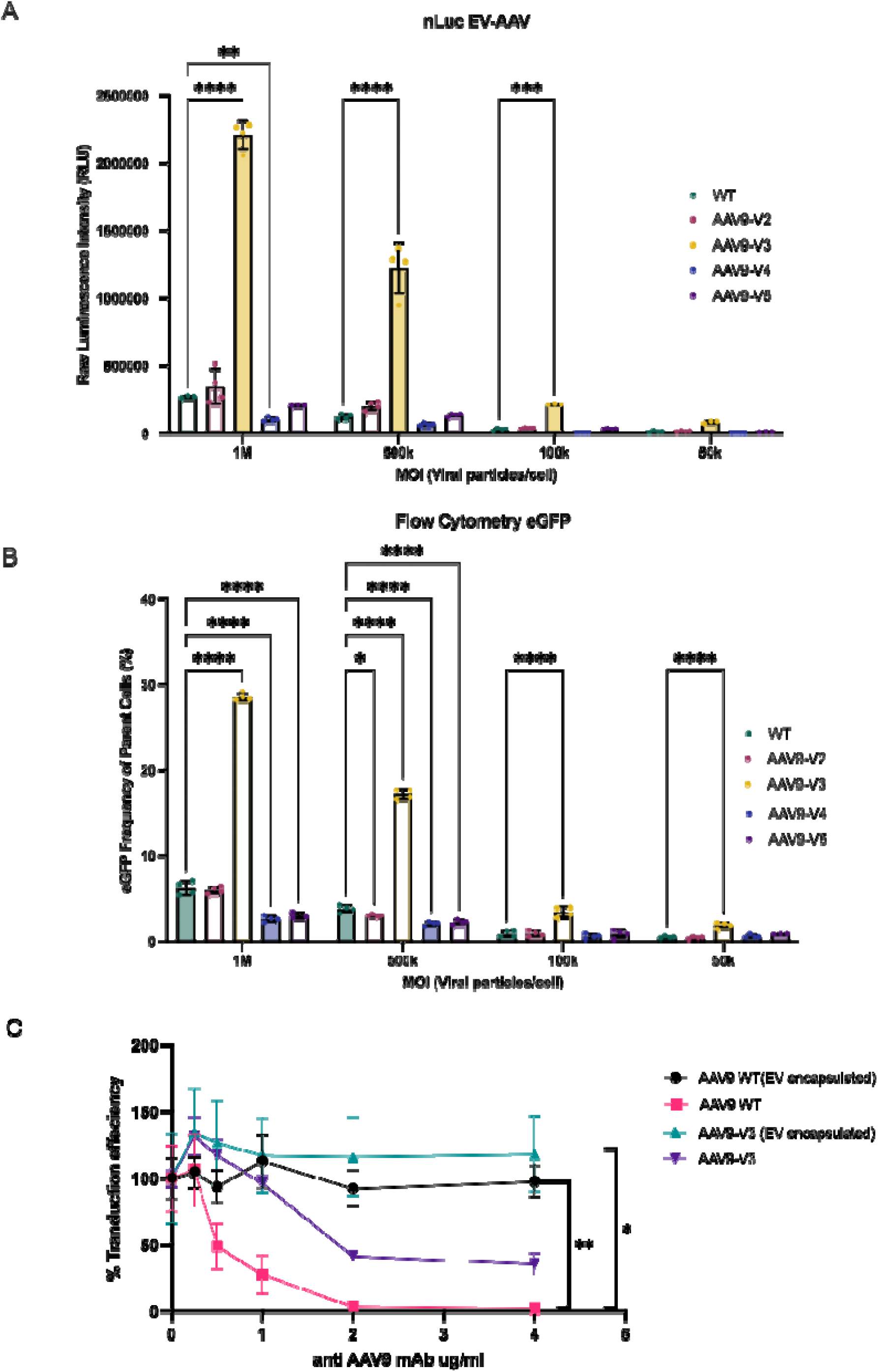
AAV9-V3 shows improved *in vitro* transduction in comparison to AAV9 wildtype. (A) Bulk transduction efficiency quantified using the NanoLuc–eGFP fusion reporter encoded in all vectors. HeLa cells were transduced with non-encapsulated WT AAV9 or engineered variants (V1–V5) across a range of MOIs. Total NanoLuc luminescence was measured from cell lysates 24 h post-transduction. Increased luminescence reflects higher intracellular delivery and reporter expression. (B) Single-cell transduction efficiency determined by the percentage of eGFP-positive HeLa cells measured by flow cytometry. As in (A), these data reflect non-encapsulated AAV preparations. The NanoLuc–eGFP fusion enables both bulk and single-cell readouts from the same transgene cassette. (C) Neutralisation assay evaluating susceptibility of WT AAV9 and AAV9-V3 to escalating concentrations of anti-AAV9 neutralising antibodies. Both free AAV and EV-encapsulated AAV (EV-AAV) preparations were tested, as indicated in the legend. EV encapsulation consistently improved neutralisation tolerance, with EV-AAV9-V3 showing the greatest resistance and maintaining robust transduction even at the highest antibody titres. Transduction was quantified as the percentage of eGFP-positive cells. All values represent mean ± SEM from n = 3 biological replicates. Statistical analysis was performed using two-way ANOVA with Dunnett’s multiple comparisons test relative to WT AAV9. **P < 0.01; *P < 0.001.

Encapsulation of vectors within extracellular vesicles increased overall transduction efficiency for all capsids; however, EV-AAV9-V3 remained the most potent configuration, demonstrating that the V3 capsid retains its functional advantage when associated with EVs. EV-AAV9-V3 preparations displayed the expected EV characteristics: Western blotting confirmed the presence of canonical EV markers (CD63, CD81, and Alix), verifying successful and specific enrichment of extracellular vesicles without detectable cellular contamination (Supplementary Figure 2).

To determine whether the engineered capsids differ in susceptibility to humoral inhibition, we assessed vector function in the presence of escalating concentrations of anti-AAV9 neutralising antibodies. Wild-type AAV9 showed a sharp, dose-dependent loss of transduction, consistent with its known vulnerability to antibody-mediated neutralisation (Figure 5C). In contrast, AAV9-V3 retained significantly higher transduction at all antibody titres, exhibiting a right-shifted and markedly flatter neutralisation curve, indicative of enhanced resistance. EV encapsulation provided additional protection: EV-WT AAV9 showed partial rescue at intermediate antibody concentrations, whereas EV-AAV9-V3 displayed the strongest neutralisation tolerance, maintaining robust transduction even at the highest antibody levels tested. The combined effects of capsid engineering and EV-mediated shielding produced the greatest preservation of functional activity across the entire neutralisation range.

## Discussion

EV-AAV vectors offer a promising platform to overcome several limitations of conventional AAV-based gene therapy, including susceptibility to neutralising antibodies and inefficient cellular uptake^22–24^. However, endogenous loading of AAV into EVs is typically inefficient, with only 0.5% to 12% of total AAV genomes encapsulated in EVs, depending on AAV serotype^13^, limiting the scalability of this approach. With AAV-based gene therapy production already being resource-intensive, for EV-AAV therapy to become clinically viable, we would need to reach encapsulation efficiencies in the higher double-digit range, to reach AAV genome titres comparable to conventional AAV-based therapies. While published AAV9 wild-type titres lie around 1.75x10^10^ viral genomes per millilitre in EV fractions ^13^, we were able to increase this to approximately 3x10^11^ viral genomes per millilitre, demonstrating that clinical translation might be within reach using our engineering approach.

In this study, we applied a combined rational design and directed evolution strategy to engineer AAV9 capsids with enhanced EV incorporation and functionality. By inserting peptide motifs derived from viral late domains and EV-sorting signals into a surface-exposed loop of the AAV9 capsid with random linkers of varying length on either end, we generated a diverse capsid library and used iterative selection and enrichment analysis to identify variants with superior EV association. Our top five variants demonstrated up to a threefold increase in viral genome packaging into EVs compared to wild-type controls. Importantly, increased EV loading was not associated with changes in vesicle size or abundance, suggesting that enhanced association resulted from specific interactions between capsid motifs and host EV biogenesis pathways rather than global changes in EV production.

The identification of functional motifs within enriched variants supports the mechanistic rationale behind our engineering strategy. For example, the SVPPHHAYVPTG variant contains a PPXnY-like motif commonly used by enveloped viruses to engage the ESCRT machinery via NEDD4 family ubiquitin ligases, while LSDRKIERQGNVF shares similarity with the KFERQ motif known to promote lysosomal targeting via LAMP2A and has been implicated in protein sorting into EVs. These results align with previous studies, showing that certain sorting signals or membrane-targeting sequences can enhance capsid association with lipid bilayers or facilitate budding into vesicles. LRGDVGGP was the third most enriched insertion, it does not contain any Late domain motifs but does contain the residues RGD. Incorporation of an RGD motif into the capsid may promote its loading into EVs by facilitating interactions with integrins, which are abundantly expressed on the surface of EV-secreting cells^25^. RGD–integrin binding can trigger endocytosis and influence vesicular trafficking pathways, including sorting into multivesicular bodies (MVBs), which give rise to exosomes^26^. Through this mechanism, RGD-modified capsids may be preferentially directed into EVs during membrane budding or MVB fusion, thereby enhancing their encapsulation efficiency.

Biophysical assays confirmed that engineered capsid variants not only increased total EV-associated genome counts but also promoted true encapsulation within EVs, as measured by protection from antibody binding in detergent-sensitive ELISAs. A 100 nm extracellular vesicle can theoretically fit up to ∼47 AAV particles under ideal packing conditions, assuming the EV is otherwise empty and packing efficiency is maximised. However, due to geometric constraints inside a spherical container, realistic packing is closer to 50–60%, allowing for ∼32–39 AAVs at most. Future work could involve Cryo-EM microscopy experiments to validate packaging density.

*In vitro* neutralization assays demonstrated that engineered EV-AAV9s were significantly more resistant to pooled human IVIG compared to both free AAV9 and wildtype EV-AAV9.

Our screening approach also highlights the value of integrating statistical modelling into library design and analysis, particularly when it is infeasible to generate technical replicates. Statistical modelling allows for providing an estimate for the variability of measurements based on random noise; knowing the uncertainty of these measurements is crucial for establishing an effective assay that distinguishes signal from noise, and for effective identification of candidate/s to carry forward for further analyses. Here, we applied statistical modelling to define a quality cutoff score and motif analysis to average the enrichment score of motifs over multiple sequences. This strategy offers a broadly applicable workflow for optimising capsid libraries or other sequencing-based directed evolution screens, allowing for maximised throughput. This could be done by building on the statistical model to identify ideal parameters (e.g., reducing subsampling during NGS or when extracting variants for subsequent steps), reducing cross-packaging, or lowering the noise threshold by generating technical replicates.

More broadly, our research has highlighted the need for designing assays with optimal trade-off between throughput and robustness. Such well-balanced assays would maximise the number of variants screened, while ensuring that each design screened is measured with significant signal-to-noise ratio. This will become even more important when designing screens to generate data for machine learning approaches. Similarly, knowledge about the signal-to-noise ratio will allow us to accurately identify candidates to carry forward for future analysis, which is particularly important as some may fail downstream tests, for example, *in vitro* transduction efficiency evaluation (see Figure 5).

While the directed evolution process successfully enriched for functional capsid variants, opportunities remain to further optimise and expand this approach. For example, library coverage in the current screen was influenced by technical factors such as sequencing depth and insert size bias, which may have limited the representation of certain high-performing variants. Nonetheless, the inclusion of longer variants in the secondary screen helped broaden the landscape of tested candidates. Building on this platform, engineered capsid variants can be paired with producer cell modifications—for instance, overexpressing LAMP2A in cells generating LSDRKIERQGNVF-AAV9—to further boost EV-AAV yields^27^. Additionally, enrichment data from this screen provides a valuable training set for machine learning models aimed at predicting novel capsid sequences with improved performance, even beyond those captured experimentally.

In summary, this study demonstrates that rational capsid engineering combined with high-throughput screening represents a promising strategy to move the EV-AAV technology towards clinical translation. As pre-existing immunity to AAV capsids remains a major barrier to broad patient eligibility and therapeutic efficacy, strategies that enable immune evasion without compromising vector potency are urgently needed. Engineered EV-AAVs with improved capsid encapsulation offer a promising solution, potentially expanding access to AAV gene therapies for patients with high NAb titres. Furthermore, the scalable and modular nature of this platform opens avenues for tailoring EV-AAVs to specific tissues or diseases, accelerating the development of next-generation vectors with enhanced safety, targeting precision, and therapeutic durability.

## Material and Methods

### Cell Culture

HEK293T (ATCC CRL-11268) cells were cultured in Dulbecco’s Modified Eagle Medium (DMEM; Life Technologies, Cat. #11995065) supplemented with 10% heat-inactivated fetal bovine serum (FBS; Life Technologies, Cat. #26140079) and 1× Antibiotic-Antimycotic (Sigma-Aldrich, Cat. #A5955). Cultures were maintained at 37 °C in a humidified incubator with 5% CO. Cells were routinely screened for mycoplasma contamination using a PCR Mycoplasma Test Kit I/C (PromoKine, Cat. #PK-CA91-1024-02). Cell confluency was monitored using brightfield microscopy and maintained between 50–80% for transfection and virus production workflows.

### AAV Library Design and Cloning

Peptide libraries were designed to introduce either known L domain motifs or fully randomized sequences into the VR-IV loop of the AAV9 capsid (VP1 residue 453). Inserts were flanked by glycine-serine linkers of varying lengths (2, 4, or 8 residues), allowing systematic evaluation of motif accessibility and EV engagement. Central regions included PTAP, YPXnL, PPXY, PLPPV, LXXL, or KFERQ motifs, alongside randomized controls of equivalent length. All sequences were codon-optimized and flanked with BsmBI-compatible overhangs. Single-stranded oligos (Integrated DNA Technologies) were converted to double-stranded DNA via overnight extension with Klenow fragment (exo–, NEB, Cat. #M0212S) in a 25 μL reaction containing 1 μL oligo (100 μM), 0.25 μL Klenow, 2 μL reverse primer (10 μM), 1 μL dNTPs (10 mM each), and 20.75 μL RNase/DNase-free water. Reaction products were purified using the Monarch PCR Cleanup Kit (NEB, Cat. #T1030S) and eluted in 20 μL water.

RepCap2/9-ITR plasmids were digested with BsmBI-v2 (NEB, Cat. #R0739S), gel-purified, and assembled with dsDNA inserts using Gibson Assembly Master Mix (NEB, Cat. #E2611). Reactions were incubated at 50 °C for 1 hour and purified again prior to electroporation into NEB 10-beta electrocompetent E. coli (Cat. #C3020K). Cells were recovered in SOC media and plated on LB-agar with 100 μg/mL ampicillin. After electroporation, SOC medium was added within 30 s, and bacteria were incubated in a 2 mL tube at 37°C with shaking. Around 40 µL of bacterial solution was plated on ampicillin plates and incubated upside down at 37°C overnight. Transformation efficiency was estimated by plating a 1000-fold dilution and counting colonies.

Bacterial colonies were scraped from ampicillin plates using 2–5 mL of cold LB, transferred to 50 mL Falcon tubes, and pelleted (4000 x g, 15–20 min, 4°C). Supernatants were discarded, and pellets (∼0.45 g) were processed using Qiagen or Macherey-Nagel Maxiprep kits per manufacturer protocols. Eluates from multiple columns were combined if necessary. Samples were sent for Sanger sequencing for confirmation. Final plasmid pools were stored at –20 °C.

### AAV Production and Directed Evolution Screening

HEK293T cells were seeded at 8×10 cells per 150 cm² dish (Starlab) 24 hours prior to transfection. At 70–80% confluency, cells were transfected using linear 25 kDa PEI (Polysciences, Cat. #23966-2) at a DNA:PEI ratio of 1:2.5 (w/w). For library screening, each dish received a total of 50 μg DNA: 25 μg helper plasmid (encoding adenoviral helper genes), 13.4 μg RepCap2/9-ITR library plasmid, and 11.6 μg transgene vector encoding GFP. Transfection complexes were incubated in Opti-MEM (Gibco, Cat. #31985070) for 20 minutes prior to dropwise addition to culture media. After 4–6 hours, media was replaced with fresh OptiMEM (Gibco, Cat. #31985070). After 48 hours, supernatant was harvested and clarified by centrifugation at 400×g for 5 minutes followed by 4,000×g for 20 minutes. For library evolution, DNA was extracted from purified EV-AAV9 samples, amplified by PCR using insert-specific primers, and re-cloned into fresh RepCap2/9-ITR backbones as described above. This process was repeated for three rounds, enriching capsids with improved EV incorporation.

### Extracellular Vesicle Isolation via Iodixanol Gradient

Clarified media was concentrated to 500 μL using tangential flow filtration (TFF) and Amicon Ultra-15 spin filters (Millipore, 100 kDa MWCO, Cat. #UFC910024). EVs and associated AAVs were purified using a discontinuous iodixanol gradient prepared in 14 × 89 mm OptiSeal tubes (Beckman Coulter, Cat. #361625): 2 mL 60%, 2 mL 40%, 2 mL 25%, 1 mL each of 18%, 16%, 14%, 12%, and 6% iodixanol (OptiPrep, Sigma-Aldrich, Cat. #D1556). 500 μL concentrated media was layered gently on top and ultracentrifuged at 28,500 rpm (Beckman SW41 Ti rotor) for 16 hours at 4 °C. Fractions (1 mL each) were collected from the top. EVs were typically recovered from fractions 4–6, while AAV particles peaked in fractions 9–12. Iodixanol was removed by washing four times with sterile PBS using 100 kDa spin filters. Final concentrates were stored at 4 °C short-term or –80 °C for long-term storage.

### Viral Genome Titration (qPCR and ddPCR)

5 μL of each sample was DNase-treated (Promega, Cat. #M6101) at 37 °C for 30 minutes. Reactions were stopped with 2 μL DNase stop solution, followed by incubation at 65 °C for 10 minutes. To release genomes, samples were digested with proteinase K (Roche, Cat. #03115879001) in 100 μL lysis buffer (0.1 M EDTA, 0.5% SDS) and incubated at 50 °C for 1 hour. After dilution (1:300), samples were analyzed by qPCR using Fast SYBR Green Master Mix (Applied Biosystems, Cat. #4385612) with primers against the transgene. Reactions (20 μL) included 100 nM forward/reverse primers and 3 μL template. Standard curves were generated from linearized plasmid DNA ranging from 10³–10 copies. Digital droplet PCR (ddPCR) was performed using the QX200 Droplet Digital PCR System (Bio-Rad) as needed to confirm genome copy numbers for in vivo dosing. Droplets were generated from diluted DNA and analyzed using QuantaSoft software.

### Statistical Model

We modelled the synthesis of the fully random and semi-random variant designs by randomly drawing the copy number per theoretically possible variant from a binomial distribution, using the Python library numpy. For the 20- and 21-mer semi-random and fully random variants, the probability of obtaining more than two copies of a particular variant was negligible. Thus, we assumed that there would be as many variants synthesised as there were molecules, with each variant having a copy number of 1. We numerically approximated distributions with high variant counts by modelling the copy number per variant for a fraction of the variant pool, transforming the copy number per variant array into a frequency per copy number array, and extrapolating this frequency per copy number array to the entire population. Transformation of plasmids into bacterial cells was modelled by drawing from a binomial distribution. While the transformation step theoretically involves a random draw without replacement, simulating this using the hypergeometric distribution was computationally inefficient. Therefore, since the number of molecules transformed represents less than 5% of the total number of molecules, we approximated the hypergeometric distribution using the binomial distribution^28^.

Next, we modelled the bacterial replication of the variants under the assumption that the number of copies per variant doubles as the bacterial cells double. The number of bacterial cells over time was calculated using a standard growth formula re-arranged below^29^, where r represents the growth rate, N_t1_ represents the number of bacteria at time point t1 and N_t1_ represents the initial number of bacteria. To derive the mean growth rate, we used data provided by the manufacturer of the competent cells. Since growth rates vary between bacterial cells, we drew the growth rate from a Gaussian distribution.

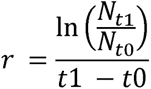

We subsequently performed two more rounds of random draws with replacement using the binomial distribution to model the isolation of plasmids from bacteria and the retrieval of high-quality reads from the variants after sequencing and bioinformatic processing. To simulate random enrichment, we drew twice from the pre-sequencing distribution and calculated enrichment scores by dividing the copy number per variant from distribution 1 by the copy number per variant from distribution 2.

### Next-Generation Sequencing and Variant Enrichment Analysis

The variable insert region was amplified using high-fidelity polymerase (KAPA HiFi, Roche) and primers flanking the VR-IV loop. Amplicons were purified using Monarch kits and submitted for sequencing (Amplicon-EZ, Azenta or BGI). If fastq files were split between different lanes, we combined them (Azenta). Paired-end reads were aligned and merged using a custom Python script. Reads were processed using Cutadapt^30^ to filter based on flanking regions and trim insert sequences in an error-tolerant manner. Next, we converted fastq files into Pandas data frames and determined the length of each insert. Inserts shorter than 24 nts or longer than 63 nts were discarded. Insert frequencies were derived and normalised to the total number of read counts. We then derived enrichment scores by merging the data frames for the original library and the enriched library based on the insert sequence and dividing the normalised variant count of the original library by the variant count in the enriched library. Finally, we translated the DNA sequence of each insert into the respective protein sequence.

Variants were filtered based on an enrichment cutoff of 10. This cutoff was chosen to eliminate 99.99% of the enrichment scores that we would expect based on random chance, as defined by our statistical model. After discarding variants with an enrichment score below the cutoff, sequences containing stop codons were discarded. Next, inserts were grouped by their length (24, 27, 36, 39, 60 and 63 nts) and based on their design approach (semi-random vs. fully random). Next, amino acid sequence logos were generated to detect amino acid motifs that are enriched over multiple sequences. We then selected the top variants with enriched motifs per sequence design (semi and fully random groups with different insert lengths). The top 120 enriched variants were selected for a secondary screen, including a positive control (wild-type) and a negative control (stop codon).

### Nanoparticle Characterization by Nanoflow Cytometry

EV-AAV samples were diluted in sterile-filtered PBS and analyzed using a NanoFCM flow nanoanalyzer (NanoFCM Inc.) calibrated with silica nanoparticles of known size (Cat. #S16M-Exo). Each sample was run in triplicate, and data were analyzed using Flow NanoAnalyzer software. Particle size distributions and concentration were quantified for each EV preparation.

### Capsid Encapsulation Assay (MSD ELISA)

AAV9 capsid accessibility was assessed with and without detergent using the Meso Scale Discovery platform. Streptavidin-coated 96-well plates (MSD GOLD, Cat. #L15SA) were coated with 0.1 μg/mL biotinylated anti-AAV9 antibody (Thermo Fisher, Cat. #7103332100) for 1 hour at 21 °C. Plates were washed with 0.05% Tween-20 in PBS using a Biotek ELx405 plate washer. Samples were diluted 1:1,000 and 1:10,000 in MSD Diluent 100, with or without pre-treatment with 0.5% Triton-X-100 (Sigma). Plates were incubated for 1 hour at 700 rpm. Detection was performed using 0.5 μg/mL ADK9-1R (Progen, Cat. #610178), followed by SULFO-TAG anti-mouse secondary antibody (MSD, Cat. #R32AC-1). Plates were read using the MESO QuickPlex SQ120 instrument. Readings were normalized to the detergent-treated condition to calculate encapsulated fraction.

### Magnetic Pulldown of AAV Particles

25 μL Dynabeads MyOne Streptavidin T1 (Thermo Fisher, Cat. #65601) were washed and incubated with 2.5 μL biotinylated anti-AAV9 antibody for 30 minutes at room temperature. After washing with 0.1% BSA/PBS, samples were added and incubated overnight at 4 °C with rotation. Beads were washed 3× and resuspended in PBS for genome quantification by qPCR.

### Nanoluc quantification

HeLa cells were seeded at 2×10 cells per well in 96-well white opaque plates and transduced with WT AAV9 or engineered AAV9 variants at the indicated MOIs. For luminescence quantification, vectors carried a CMV–NanoLuc (nLuc) reporter cassette. Twenty-four hours post-transduction, cells were washed once in PBS and lysed in 20 μL Passive Lysis Buffer (Promega). Luminescence was quantified using the Nano-Glo Luciferase Assay System (Promega, N1120) according to the manufacturer’s protocol by adding 20 μL of Nano-Glo substrate to each well and reading immediately on a CLARIOstar plate reader (BMG Labtech) with a 1-second integration time. Background-subtracted relative luminescence units (RLU) were normalised to total protein content measured by BCA assay.

### Flow Cytometric Quantification of GFP-Positive Cells

For experiments in Fig. 5B and Fig. 5C, vectors encoded an enhanced GFP (eGFP) reporter under the CMV promoter. HeLa cells were harvested 24–48 hours post-transduction, washed in PBS, and resuspended in FACS buffer (2% FBS/PBS). Flow cytometry was performed on a BD LSRFortessa using a 488-nm laser and 530/30-nm emission filter. Autofluorescence gates were set using untransduced controls. A minimum of 10,000 live singlet events were recorded per sample. Data were analysed using FlowJo v10, and eGFP-positive cells were expressed as a percentage of total viable singlets.

### Neutralization Assay

Neutralising activity against AAV9 was assessed by pre-incubating AAV vectors with increasing concentrations of anti-AAV9 antibodies before cell transduction. AAV9-specific neutralising antibodies (mouse monoclonal pool or serum source as appropriate) were serially diluted in complete DMEM to generate a dose-escalation series. For all conditions, 5 × 10 vector genomes (vg) of either WT AAV9 or AAV9-V3—EV encapsulated or non-encapsulated and carrying EGFP transgene—were incubated with the antibody dilutions at 37 °C for 1 hour to allow antibody–capsid interaction. After incubation, the antibody–virus mixtures were added directly to HeLa cells (70–80% confluence) in 24-well plates. Transduction efficiency was quantified 24–48 hours later by flow cytometry for reporter expression. All conditions were performed in technical triplicates.

## Data and Code Availability

All of the raw data is available on request. We provide a generalizable Jupyter notebook for applying the statistical model to other experimental designs. To adjust the model to your specification, insert your parameters into the JSON file and run the notebook (https://github.com/marinaluchner/StochasticModeling). We further provide the Jupyter notebooks for the bioinformatic pipeline used for this paper in a GitHub repository (https://github.com/marinaluchner/BioinformaticAnalysis).

## Author Contributions

I.V. and M.L. contributed equally. I.V., M.L., and D.G. conceived and designed the study, with input from M.P.A.W. and S.J.S. M.L. developed the SMAPS statistical framework and performed computational analyses. I.V. and co-authors performed experiments and data analysis.

M.P.A.W. and D.G. supervised the work. I.V., M.L., and D.G. wrote the manuscript, with input from all authors.

## Acknowledgment

I.V. was supported by a DPhil studentship from Evox Therapeutics.

M.L. received funding from the Biotechnology and Biological Sciences Research Council (BBSRC), UK Research and Innovation (UKRI), under grant BB/T008784/1.

D.G. is supported by a Hjärnfonden Postdoctoral Fellowship and the British Heart Foundation (BHF) CureHeart programme. This work was additionally supported by the Cystic Fibrosis Foundation (CFF; HNR02540), the U.S. Department of Defense (DoD; A3R00010; W81XWH-22-1-1067), and Muscular Dystrophy UK (MDUK; HNR01740; 19GRO-PG36-0339).

## Conflict of interest

D.G is a stakeholder in Evox Therapeutic Ltd (UK), M.J A.W is a consultant and stakeholder in Evox Therapeutic Ltd (UK). The rest of the co-authors reported no conflict of interest.

## References

1. Naso, M. F., Tomkowicz, B., Perry, W. L. & Strohl, W. R. Adeno-Associated Virus (AAV) as a Vector for Gene Therapy. Biodrugs 31, 317–334 (2017).

2. Wang, D., Tai, P. W. L. & Gao, G. Adeno-Associated Virus Vector as a Platform for Gene Therapy Delivery. Nat Rev Cancer 18, 358–378 (2019).

3. Wang, J.-H., Gessler, D. J., Zhan, W., Gallagher, T. L. & Gao, G. Adeno-associated virus as a delivery vector for gene therapy of human diseases. Signal Transduct Target Ther 9, 78 (2024).

4. Loo, J. C. M. van der & Wright, J. F. Progress and Challenges in Viral Vector Manufacturing. Hum Mol Genet 25, R42–R52 (2015).

5. Hamilton, B. A. & Wright, J. F. Challenges Posed by Immune Responses to AAV Vectors: Addressing Root Causes. Front Immunol 12, (2021).

6. Riyad, J. M. & Weber, T. Intracellular trafficking of adeno-associated virus (AAV) vectors: challenges and future directions. Gene Ther 28, 683–696 (2021).

7. 7. Raposo, G. & Stoorvogel, W. Extracellular vesicles: Exosomes, microvesicles, and friends. Journal of Cell Biology vol. 200 373–383 Preprint at 10.1083/jcb.201211138 (2013).

8. Maguire, C. A. Accessorizing viral vectors with extracellular vesicles for enhanced performance. Molecular Therapy 31, 1204–1206 (2023).

9. Deshetty, U. M. Gene Therapy for the Heart: Encapsulated Viruses to the Rescue. Extracell Vesicles Circ Nucl Acids (2024) doi:10.20517/evcna.2023.70.

10. 10. György, B. & Maguire, C. A. Extracellular vesicles: nature’s nanoparticles for improving gene transfer with adeno-associated virus vectors. Wiley Interdisciplinary Reviews: Nanomedicine and Nanobiotechnology vol. 10 Preprint at 10.1002/wnan.1488 (2018).

11. György, B., Fitzpatrick, Z., Crommentuijn, M. H. W., Mu, D. & Maguire, C. A. Naturally enveloped AAV vectors for shielding neutralizing antibodies and robust gene delivery invivo. Biomaterials 35, 7598–7609 (2014).

12. Cheng, M. et al. Neutralizing Antibody Evasion and Transduction with Purified Extracellular Vesicle-Enveloped Adeno-Associated Virus Vectors. Hum Gene Ther 32, 1457–1470 (2021).

13. Cheng, M. et al. Probing aspects of extracellular vesicle associated AAV allows increased vector yield and insight into its transduction and immune-evasive properties. Mol Ther Methods Clin Dev 33, (2025).

14. Ferreira, J. V. et al. LAMP2A regulates the loading of proteins into exosomes. Sci Adv 8, 1140 (2022).

15. Freed, E. O. Viral Late Domains. J Virol 76, 4679–4687 (2002).

16. Welker, L., Paillart, J. C. & Bernacchi, S. Importance of viral late domains in budding and release of enveloped rna viruses. Viruses vol. 13 Preprint at 10.3390/v13081559 (2021).

17. Hoffmann, M. D., Gallant, J. P., LeBeau, A. M. & Schmidt, D. Unlocking precision gene therapy: harnessing AAV tropism with nanobody swapping at capsid hotspots. NAR Molecular Medicine 1, (2024).

18. Tabebordbar, M. et al. Directed evolution of a family of AAV capsid variants enabling potent muscle-directed gene delivery across species. Cell 184, 4919–4938.e22 (2021).

19. Maheshri, N., Koerber, J. T., Kaspar, B. K. & Schaffer, D. V. Directed evolution of adeno-associated virus yields enhanced gene delivery vectors. Nat Biotechnol 24, 198–204 (2006).

20. Arnold, F. H. Design by Directed Evolution. Acc Chem Res 31, 125–131 (1998).

21. Gečys, D. et al. Internalisation of RGD-Engineered Extracellular Vesicles by Glioblastoma Cells. Biology (Basel) 11, (2022).

22. Hudry, E. et al. Exosome-associated AAV vector as a robust and convenient neuroscience tool. Gene Ther 23, 380–392 (2016).

23. Wassmer, S. J., Carvalho, L. S., György, B., Vandenberghe, L. H. & Maguire, C. A. Exosome-associated AAV2 vector mediates robust gene delivery into the murine retina upon intravitreal injection. Sci Rep 7, (2017).

24. György, B. et al. Rescue of Hearing by Gene Delivery to Inner-Ear Hair Cells Using Exosome-Associated AAV. Molecular Therapy 25, 379–391 (2017).

25. Ruoslahti, E. RGD and other recognition sequences for integrins. Annu Rev Cell Dev Biol 12, 697–715 (1996).

26. Feng, S., Zhang, Y., Liu, H., Liu, T. & Chen, Y. Cellular internalization of RGD-modified nanoparticles by integrin-mediated endocytosis and the effect of nanoparticle size. ACS Nano 5, 6727–6737 (2010).

27. Ferreira, J. V. et al. LAMP2A regulates the loading of proteins into exosomes. Sci Adv 8, (2022).

28. Weiss, N. A. Introductory Statistics. (Pearson Education Limited, Harlow, 2017).

29. Razo-Mejia, M., Mani, M. & Petrov, D. Bayesian inference of relative fitness on high-throughput pooled competition assays. PLoS Comput Biol 20, e1011937-(2024).

30. Martin, M. Cutadapt removes adapter sequences from high-throughput sequencing reads. EMBnet J 17, 10–12 (2011).

31. Dolnik, O. et al. Interaction With Tsg101 Is Necessary for the Efficient Transport and Release of Nucleocapsids in Marburg Virus-Infected Cells. PLoS Pathog (2014) doi:10.1371/journal.ppat.1004463.

32. Kenney, S. P., Wentworth, J. L., Heffron, C. L. & Meng, X. Replacement of the Hepatitis E Virus ORF3 Protein PxxP Motif With Heterologous Late Domain Motifs Affects Virus Release via Interaction With TSG101. Virology (2015) doi:10.1016/j.virol.2015.09.012.

33. Chen, X. ALIX and TSG101 Are Essential for Cellular Entry and Replication of Two Porcine Alphacoronaviruses. PLoS Pathog (2024) doi:10.1371/journal.ppat.1012103.

34. González López, O., et al. Redundant Late Domain Functions of Tandem VP2 YPX _3_ L Motifs in Nonlytic Cellular Egress of Quasi-Enveloped Hepatitis a Virus. J Virol (2018) doi:10.1128/jvi.01308-18.

35. Maaroufi, H. SARS-CoV-2 Encodes a PPxY Late Domain Motif Known to Enhance Budding and Spread in Enveloped RNA Viruses. (2020) doi:10.1101/2020.04.20.052217.

36. Shirasaki, T. et al. Nonlytic Cellular Release of Hepatitis a Virus Requires Dual Capsid Recruitment of the ESCRT-associated Bro1 Domain Proteins HD-PTP and ALIX. PLoS Pathog (2022) doi:10.1371/journal.ppat.1010543.

37. Vecchio, C. Del et al. Alix-Mediated Rescue of Feline Immunodeficiency Virus Budding Differs from That Observed with Human Immunodeficiency Virus. J Virol 94, 10.1128/jvi.02019-19 (2020).

38. Coren, L. V, Nagashima, K. & Ott, D. E. A PLPPV sequence in the p8 region of Gag provides late domain function for mouse mammary tumor virus. Virology 535, 272–278 (2019).

39. Pei, Y. et al. Mutation of Phenylalanine 23 of Newcastle Disease Virus Matrix Protein Inhibits Virus Release by Disrupting the Interaction between the FPIV L-Domain and Charged Multivesicular Body Protein 4B. Microbiol Spectr 11, e04116–22 (2023).

40. Matsuura, R. et al. Three YXXL Sequences of a Bovine Leukemia Virus Transmembrane Protein are Independently Required for Fusion Activity by Controlling Expression on the Cell Membrane. Viruses 11, (2019).

