## Supplementary Figures for "Engineered AAV Capsids with Enhanced Extracellular Vesicle Loading via Rational Design and Directed Evolution"

### **Supplementary Information**

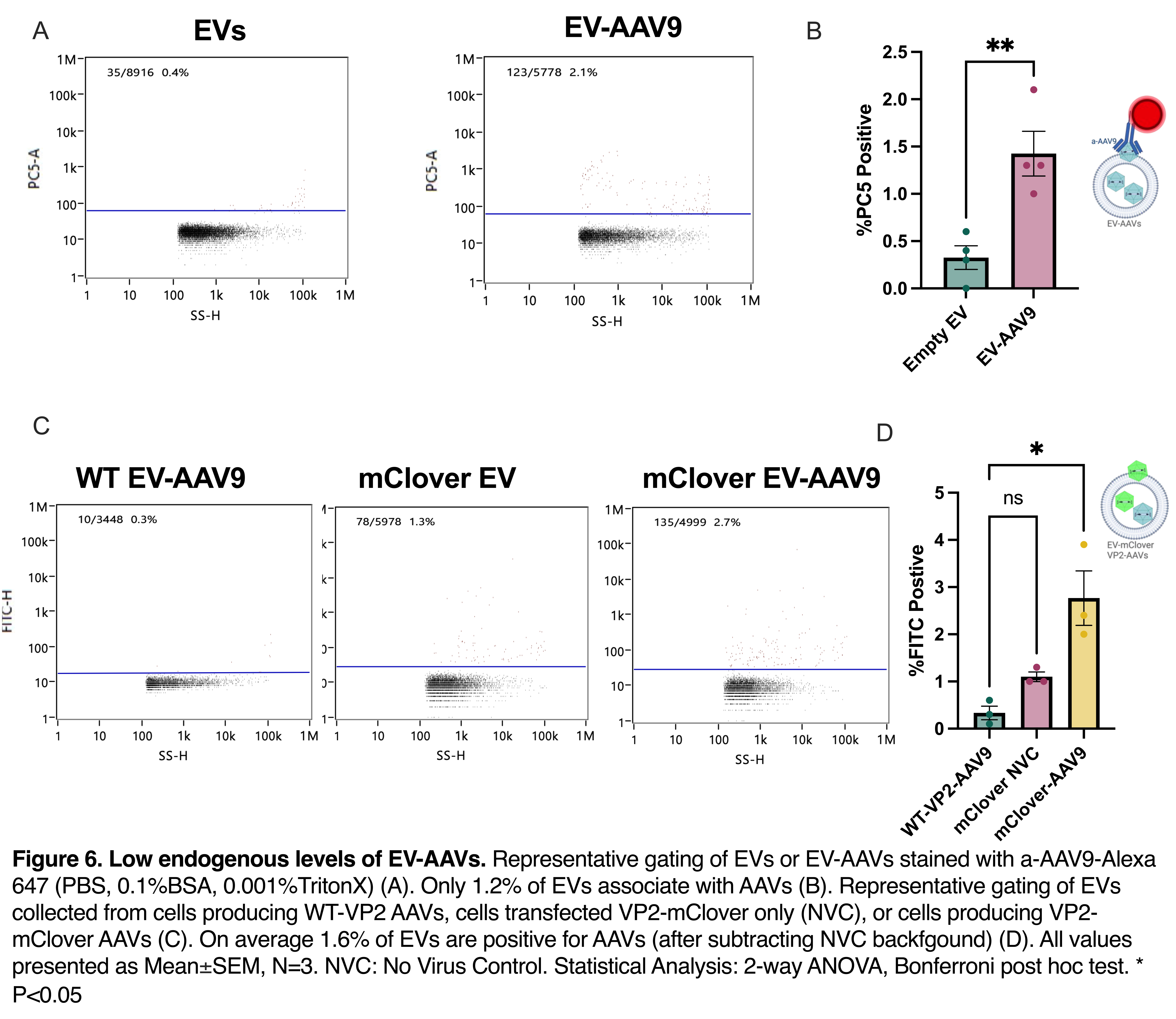

Supplementary Figure 1. Low endogenous levels of EV-AAV9s.

Representative gating of EV or EV-AAV9 stained with anti-AAV9-Alexa 647 (PBS, 0.1% BSA, 0.001% Triton-X-100) (A). Only 1.2% of EVs are associated with AAVs (B). Representative gating of EVs collected from cells producing WT-VP2 AAVs, cells transfected VP2-mClover only (NVC: No virus control), or cells producing VP2-mClover AAVs (C). On average, 1.6% of EVs are positive for AAVs (after subtracting NVC background) (D). All values are presented as Mean±SEM. NVC: No Virus Control. Statistical Analysis: F-test for equality of variances: *F*(3, 3) = 3.560, *p* = 0.3248. Unpaired two-tailed *t*-test: *t*(6) = 4.121, *p* = 0.0062 (B). Ordinary one-way ANOVA detected a significant difference among groups (*F*(2, 6) = 12.70, *p* = 0.0070). Variances were similar based on Brown-Forsythe (*p* = 0.3758). Tukey’s test showed significant differences between WT-VP2-AAV9 and mClover-AAV9 (*p* = 0.0063), and between mClover NVC and mClover-AAV9 (*p* = 0.0344) (D).

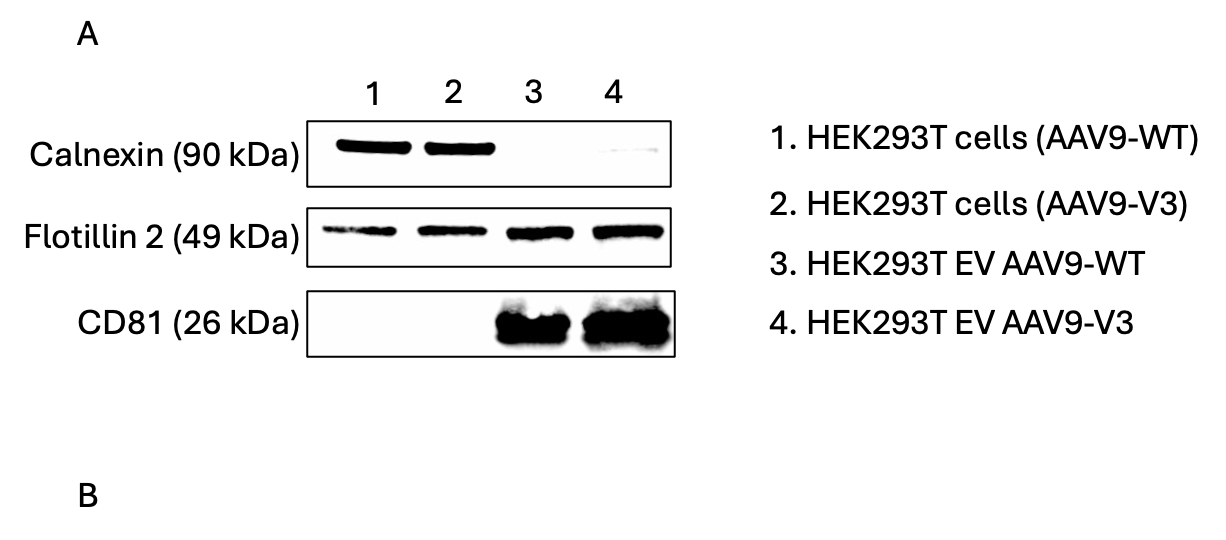

Supplementary Figure 2. Characterisation of EV-Encapsulated AAV9 WT and V3 Variants.

Western blot analysis confirming the enrichment of canonical EV markers and absence of apoptotic vesicle markers in isolated EV preparations. EV samples derived from each AAV9 capsid condition show strong expression of Flotillin2 and CD81.

**Table S1. Amino Acid Motifs Associated with Enveloped Virus Egress, Late Domain Activity, or Protein Targeting to Extracellular Vesicles.**

| Amino Acid Motif | Virus Type | Example Viruses or Proteins | Known Interacting Partners |
| --- | --- | --- | --- |
| PT/SAP | Retroviruses, Paramyxoviruses | HIV-1, MLV (Murine Leukemia Virus), Human Parainfluenza Virus (HPIV), Hepatitis C Virus | TSG101 (Tumor Susceptibility Gene 101)^31–33^ |
| YPXnL | Retroviruses, Picornaviruses | HIV-1, SIV (Simian Immunodeficiency Virus), Hepatitis A Virus (HAV) | ALIX (PDCD6IP), ESCRT-III components^342^ |
| PPXY | Arenaviruses, Coronaviruses | Lassa Virus, Sars-CoV-2 | NEDD4 family ubiquitin ligases (e.g., ITCH, WWP1)^35^ |
| PxxP | Hepatitis E Virus | Hepatitis E Virus | TSG101^32,36^ |
| LXXL | Lentivirus | Feline immunodeficiency Virus (FIV) | ALIX^37^ |
| PPLPV | Retroviruses | Mouse mammary tumor virus (MMTV) | Possibly interacts with ESCRT components ^38^ |
| FPIV | Paramyxoviruses | Newcastle disease virus (NDV) | Charged multivesicular body protein 4 (CHMP4)^39^ |
| YXXL | Deltaretrovirus | Bovine leukemia virus (BLV) | AP-2 complex, Clathrin-mediated endocytosis components^40^ |
| KFERQ | Various Proteins | HIF1A | LAMP2A and HSC70^14^ |

This table summarizes amino acid motifs with functional roles in enveloped virus egress, late domain interactions, or the targeting of proteins into EVs. The first column lists the amino acid sequences, the second column provides examples of viruses or proteins where these sequences are found, and the third column identifies potential interacting partners that facilitate cargo loading into EVs. Abbreviations for amino acids: P = Proline, T = Threonine, S = Serine, A = Alanine, Y = Tyrosine, X = Any amino acid, L = Leucine, V = Valine, F = Phenylalanine, K = Lysine, E = Glutamate, R = Arginine, Q = Glutamine.

**Table S2. Enrichment scores of top five enriched variants**

| **Variant Name** | **Amino acid Sequence** | **Enrichment score** |
| --- | --- | --- |
| AAV9-V1 | SVPPHHAYVPTG | 3.85642161 |
| AAV9-V2 | LSGGLDILPKLC | 3.71284728 |
| AAV9-V3 | LRGDVGGP | 3.69158323 |
| AAV9-V4 | GRPLPPVTG | 3.42975605 |
| AAV9-V5 | LSDRKIERQGNVF | 3.36203317 |

This table lists the average enrichment score of each of the top 5 enriched variants, across 3 independent screens. Abbreviations for amino acids: P = Proline, T = Threonine, S = Serine, A = Alanine, Y = Tyrosine, X = Any amino acid, L = Leucine, V = Valine, F = Phenylalanine, K = Lysine, E = Glutamate, R = Arginine, Q = Glutamine.
